# Interactions of *Plasmodium falciparum* ETRAMP14.1 with PfEMP1, translocon components and other ETRAMP members at the interface of host-parasite

**DOI:** 10.1101/775650

**Authors:** Kirthana Mysore Vasudevarao Sindhe, Ananya Ray, Sanjeev Kumar, Namita Surolia

## Abstract

The Early Transcribed Membrane Proteins (ETRAMPs) belong to a multigene family which are conserved, are specific to *Plasmodium* species, and abundantly present in parasitophorous vacuolar membrane (PVM). The functions of the members of this family are poorly understood. *Pf*ETRAMP14.1 (PF3D7_1401400) is a member of this family, present only in *Plasmodium falciparum*. In this study, we report the potential interacting partners of *Pf*ETRAMP14.1 by using immunoprecipitation (IP) LC-MS/MS as well as protein-interaction network reconstructed on *in vivo* array analyses of severe malaria inflicted patients from malaria endemic Indian regions. We find *Pf*ETRAMP14.1 to be the most highly transcribed gene in severe infection. Our studies with western blot analysis and Immuno-flurorescence show that *Pf*ETRAMP14.1 is expressed at PVM during all the intraerythrocytic stages of *P.falciparum* with maximum expression at early trophozoite stage. Further, our results reveal interactions of ETRAMP14.1 with *Plasmodium falciparum* erythrocyte membrane protein 1(PfEMP1), thioredoxin (TRX2), export protein 2 (EXP2), heat shock protein 70-1 (Hsp70-1) and some of the ETRAMP family members. We propose that ETRAMP14.1 helps trafficking of PfEMP1 to the host RBC surface in conjunction with translocon machinery and the chaperone HSP 70-1.

## Introduction

*Plasmodium falciparum*, the most virulent human malaria parasite is responsible for approximately 214 million new cases and estimated 438000 deaths world-wide per year1. *Falciparum* infection presents widely different clinical outcomes varying from flu like symptoms to coma and death^2,3,4^. The parasite evades the host immune response by residing within a vacuole surrounded by two-membranes; the parasite plasma membrane (PPM) and the parasitophorous vacuolar membrane (PVM). PVM is formed during invasion of the host RBC which involves various molecular steps that allow the parasite to enter the host RBC and be encapsulated within the two membranes. PVM is the only mode of contact of the parasite with the host cell and hence to gain access to the host RBC, parasite must remodel the host to import nutrients, dispose waste products, and export proteins across it. PVM is thus considered as the primary site of host-parasite interactions, playing a key role in survival and propagation of the intracellular parasites. PVM resident proteins considered as ideal targets for novel anti-malarial intervention strategies include the exported protein 1 (EXP1), components of *Plasmodium* translocon of exported proteins (PTEX) and some of the ETRAMP family members^5,6,7,8^. The functions of majority of these proteins remain unknown. ETRAMPs are single pass transmembrane proteins and were first identified using a ring stage-specific cDNA library^9^, which further led to the identification of other ETRAMPs in *Plasmodium falciparum*^10^. A total of fourteen ETRAMPS are reported to be present in *Plasmodium falciparum*. Six of the members, belong to a coherent group transcribed exclusively in the ring stage which are replaced by next set of ETRAMPs expressed during the transition of the parasite from ring to trophozoite and to schizont stages, thus exhibiting a developmental regulation, fulfilling the needs of various stages. It is reported that some members of this family translocate beyond the PVM to host erythrocytes during intra erythrocytic development of the parasite^10^. All the members of ETRAMP family are encoded by a single exon and have a conserved structure with an N-terminal signal peptide, a short lysine rich domain followed by a transmembranedomain (TM), and a highly charged variable C-terminal domain. The presence of ETRAMP homologs in other *Plasmodium* species indicates that this family is involved in tasks common to all *Plasmodia*^11^.

ETRAMPs are unevenly distributed as complexes of oligomeric arrays in PVM and display dynamic change with parasite development^11^.Though, ETRAMPs were identified a decade ago, the functions of this family still remain unknown. However, many studies have proposed functions for ETRAMPs based on their location in PVM such as, junction formation between tubovesicular network (TVN) and the RBC membrane to obtain nutrients, vesicular transport processes at PVM or stabilizing PVM^10^. A *P. berghei* ETRAMP family member, SEP2 which is abundantly expressed in gametocytes as well as in mosquito and liver stages is important in gliding motility of sporozoites thereby accounting as a promising transmission-blocking candidate^12^. Furthermore, *P.yoelii* and *P. berghei*UIS genes, UIS4 and UIS3, specifically expressed in sporozoites, are ETRAMP orthologs. UIS3, the predicted ortholog of ETRAMP13 binds to host hepatocyte liver fatty-acid binding protein (L-FABP) and this interaction could be important for parasite growth and appears to be conserved in *P. falciparum*^13,14^. Knockout of UIS4 gene, the predicted ortholog of ETRAMP10.3 could not complete the liver stage development in *P.yoelii* indicating that *Pf*ETRAMP 10.3 is essential for blood stage survival15. Further, ETRAMP5 expressed in PVM of liver stages specifically interacts with human apolipoproteins Apo1, ApoB and ApoE in the yeast two-hybrid assay16 suggesting that ETRAMPs may be vital in direct host-parasite interactions and play significant role in parasite survival.

Several major advances have been made in the recent past that have significantly advanced our understanding of protein export processesindicating that the fate of proteins which are destined for export is already decided within the parasite’s endoplasmic reticulum and involves a pentameric motif (RxLxE/Q/D) called the ‘PEXEL’ motif, which is recognized and cleaved by the aspartic protease plasmepsinV^17,18,19,20,21,22^.The protein export machinery – the *Plasmodium* Translocon of Exported proteins (PTEX) is responsible for the passage of proteins across the PVM^23^ to the host RBC cytosol. PTEX comprises of heat shock protein HSP101, (a ClpA/B-like ATPase from the AAA1 superfamily commonly associated with protein translocons), a novel protein PTEX150 and the parasite exported protein 2 (EXP2). EXP2 is the potential membrane-associated channel that forms the core of PTEX complex. PTEX88 and thioredoxin 2 (TRX2) are other components of the PTEX. This protein trafficking machinery serves as a common portal for proteins involved in virulence and survival of the parasite. *Plasmodium falciparum* erythrocyte membrane protein (PfEMP1) is a virulent protein, encoded by the *var* gene family, exhibiting antigenic variation to evade host immune response and is a central player in pathogenicity of *P.falciparum*. In our literature survey, we identified that interaction studies of ETRAMP membershave not been carried out till date but there are many proposals on their functions such as they may interact with structural proteins to fix PVM in the RBCs or could interact with soluble proteins in parasite cytoplasm to act as carriers to ferry proteins out of the parasites^10^. In this study, we carried out *in vivo* differential gene expression analysis of parasites isolated from infected severe and mild malaria patient’s blood samples. Interestingly we observed that one of the ETRAMP family member, ETRAMP14.1 was maximally expressed. In our endeavor to decipher the role of ETRAMP14.1, we employed co –immunoprecipitation, LC-MS/MS and Western analysis. We show that ETRAMP14.1 interacts with PfEMP1, PVM proteins, translocon components and other ETRAMP members. Importantly, the interacting partners were also found to be present in the protein-interaction network reconstructed on *in vivo* differential gene expression data obtained from parasite RNA isolated from blood samples of patients inflicted with severe malaria validating our *in vitro* results. This study for the first time provides insights into theinteractome of a ETRAMP family memberwhich can open vistas for designing new antimalarials.

## Materials and Methods

### Strains and Antibodies

For all experiments, 3D7 strain of *Plasmodium falciparum* was used. *P. falciparum* cultures were maintained as described earlier^24^. Rabbit polyclonal antibodies were raised against the full length ETRAMP14.1. Following peptides were used for raising custom antibodies Genemed, Biotechnologies Inc.) for the proteins: α-PTEX150 (PF14_O344) “KNKNSKKNNKKKSSQKDYIR”; α-Hsp101 (PF11_0175) “GERLKRKKEIEKKLNDLKEL”. Monoclonal anti-His and anti-Flag antibodies were obtained from Sigma Aldrich, St. Louis, MO. For IFA, anti-ETRAMP14.1, anti-PTEX150, anti-Hsp101 and anti-His antibodies were used in 1:400 dilution and Alexa fluor 488 secondary antibodies were used in 1:600 dilutions. For colocalization, antibodies were labeled with Alexa fluor 568 labelling kit and used after incubating with Alexa fluor 488 secondary antibodies.

### Parasite culture

3D7 strain of *Plasmodium falciparum* was maintained in continuous culture in glass vials by Candle jar method according to the method as described^24^.Parasites were grown in O+ human erythrocytes at 37 °C in RPMI 1640 medium supplemented with 10% human serum, 25 mM sodium bicarbonate and 25 mM HEPES. Growth was monitored by Giemsa staining of thin blood smears. Parasites were synchronized using sorbitol followed by washes in PBS^25^.

### Cloning, expression and purification of recombinant ETRAMP14.1 and MS based identification of the purified protein

*Pf*ETRAMP14.1 (PF14_0016) was amplified from genomic DNA of the *in vitro* parasite cultures was PCR amplified using the 5’GGGAATTCCATATGAAAGTTTACTAATATTTTATTC-3’ and reverse 5’ CCGCTCGAGAGCTGTTGTGGCTGG-3’ primers carrying NdeI and XhoI restriction sites respectively and cloned in pET 21b(+) to obtain recombinant protein with C-terminal His tag. PfETRAMP14.1His-tagged construct was transformed into *E. coli* BL21DE3 strain to over express the protein by induction with 1 mM IPTG for 4 h at 370C. Bacterial pellet was lysed by sonication in 50 mM Tris-HCl pH 7.4, 200 mM NaCl, 10% glycerol, 1mM PMSF and protease inhibitor cocktail (Roche). Supernatant containing 20mM imidazole was incubated with Ni-NTA His•Bind® Resin (Novagen) beads with end-to-end shaking for 4 hr at 4°C. The column was initially equilibrated with lysis buffer and washed with lysis buffer containing 50 mM imidazole. The recombinant His-tag fusion protein was eluted with lysis buffer containing 200mM and 300mM imidazole respectively.

### Raising of PfETRAMP14.1 antibodies

Polyclonal rabbit antibodies were raised against recombinant PfETRAMP14.1. Protein emulsion was made in complete Freund’s adjuvant (Sigma) for subcutaneous immunization in two rabbits followed by three successive boosters in incomplete adjuvant at four week intervals.

### Parasite extract

To obtain total parasite extracts, parasites were released from RBCs by lysis with saponin for 20 min, 370C and washed with 1XPBS. Parasite pellet was resuspended in PBS containing 0.5% TritonX-100 and incubated on ice for 30 min with intermittent mixing followed by sonication (30watt, 2sec pulse, 3 times) and incubation for 4 hr at 4°C. Supernatant was obtained by centrifugation at 11000xg for 30 min. Soluble fraction was obtained with 1XPBS alone during lysis, whereas insoluble fraction was obtained with PBS-0.5% TritonX-100 or 1X SDS-sample buffer.

### Real time PCR analysis

RNA was isolated from different stages of parasite such as rings (4-16h), early trophozoites (18-24h), mature trophozoites (26–36 h) and schizonts (38–46 h) using TRI reagentTM (Sigma). RNA was treated with *Dnase I* (NEB), followed by reverse transcription with M-MuLV Reverse Transcriptase (Fermentas) and Oligo dT primers (GENEI). Primers were designed for ETRAMP14.1, E14.1F (5’-TTCATTAGCATCTGCTTTAGCC-3’) and E14.1R (5’-GCACCATCGGTTTCTTCTTT-3’) and SeryltRNA, StF (5’-AAG TAG CAG GTC ATC GTG GTT-3’ and StR, (5’-TTCGGCACATTCTTCCATAA-3’) using Genscript software to amplify approximately 150bp of the gene. Analysis of melt curves was also performed to ensure specific amplification without any secondary non-specific amplicons. PCR was carried out in final reaction volume of 20 μl using iQTM SYBR Green supermix (BIO-RAD). Control PCR reactions were performed without the RT step to make sure genomic DNA was absent. Real time PCR analysis was carried out in i-Cycler (BIO-RAD) using the following reaction conditions: 95°C for 2 min, initial denaturation, 95°C for 20 sec, 52°C for 20sec, 72°C for 30 sec for 40 cycles. Fold difference in expression of mRNA at different stages were calculated by Ct method (Real-Time PCR applications guide Bio-Rad). The housekeeping gene SeryltRNA, constitutively expressed across different stages of parasite was used as normalization control (normalized with Ct values of mature trophozoites).

### Western blot analysis

Protein extracts were separated on SDS-PAGE gel and were transferred onto polyvinylidene difluoride membrane (Millipore) using transfer buffer containing27mM Tris-Cl,48mM glycine,20% methanol for 1h in a Trans-Blot semidry electroblotter (Bio-Rad). Blots were blocked overnight with 5% skim milk and incubated with primary antibodies in PBS containing 1% bovine serum albumin. Dilutions of primary antibodies were as follows: anti-ETRAMP14.1, 1/3000; anti-PTEX 150, 1/3000; anti-HSP101, 1/3000; monoclonal anti-His antibody, 1/2000; monoclonal anti-Flag antibody, 1/2000; anti-GAPDH,1/2000; PfHsp70-1,1/1000 (raised against the C terminus of HSP70-1) and monoclonal anti-PfEMP1 antibody, 1/500.Detection by enhanced chemiluminescence (ECL plus western blotting detection reagents, Thermo Scientific) was performed after incubation with appropriate HRP-coupled secondary antibodies (1:4000, Bangalore Genei).

### Indirect Immunofluorescence assay

Infected erythrocytes were labeled according to the protocol described by Tonkin *et al*.,2004. Gametocyte cultures were provided by Dr. Pradeep (NIMR, Bangalore). After fixation, primary antibody (1:500) was added and incubated for 1hr in 3% BSA/PBS. Cells were washed three times in PBS to remove excess primary antibody. Alexafluor goat anti-rabbit 488 secondary antibody (Molecular Probes) was added with 1:500 dilution in 3% BSA/PBS for 1hr. cells were washed three times in PBS and mounted in 70% glycerol with with 5 μgml-1 concentration of 4-6-diamidino-2-phenylindole (DAPI, Sigma) and cells were mounted on cover slips. For colocalization experiment, antibody was labeled with Zenon Alexa Fluor 568 IgG labeling kit according to manufacturer’s instructions and incubated for 1 hour after primary and secondary antibody incubations. Images were captured on a confocal laser scanning microscope (LSM 510 META; Carl Zeiss) and were analysed by image analysis software. No staining was obtained when IFA was done using ETRAMP14.1 pre-immune serum indicating the specificity of α-ETRAMP14.1. Specificity of α-PTEX150 and α-Hsp101 antibodies were determined by western analysis of parasite lysate. IFAof Nup100 in TritonX-100 permeabilized cells served as control for antibody access to all the membranes RBC membrane, PVM and PPM. See figure S2.

### Selective permeabilization

Infected erythrocytes were permeabilized with streptolysin O (SLO; Sigma). 20 U of SLO was added in PBS and cells incubated for 15 min at room temperature, and then cells were washed three times in PBS and fixed. Control cells were treated with PBS only without SLO or treated with Triton X-100 which permeabilizes all the membranes. All treated cells were washed and were used for IFA.

### Immunoprecipitations

Parasite were isolated from rings and early trophozoite stages and lysates were prepared in 1X PBS containing 0.5% TritonX-100 and protease inhibitors (Roche) by sonication (amplitude 30, pulse 2 second, 3cycles) on ice. Cell lysate was cleared by centrifugation at 11,000xg for 45 min at 4°C. Protein G Sepharose beads CL-4B (GE Healthcare) were pre-equilibrated with lysis buffer. The lysate supernatant was precleared by incubation with pre-equilibrated Protein-G-sepharose for 2 h at 40C. Pre-equilibrated protein G sepharose beads were incubated with anti-ETRAMP14.1 (10 µg) antibodies for 2 h at 4°C. Antibody bound protein G sepharose was incubated for 6 h with pre-cleared lysate at 4°C. The immunoprecipitated complex was washed three times with the lysis buffer and run on SDS-PAGE. The presence of ETRAMP14.1 was confirmed by Western analysis in the immunoprecipitate. A number of bands from the gel were excised and analysed by LC-MS/MS.

### LC-MS/MS Analysis

Bands of interest in silver stained gel were rinsedand in gel digestion was performed. Peptides extracted were analyzed on LTQ-OrbitrapVelos mass spectrometer (Thermo Scientific, Bremen, GmbH) interfaced with Easy nano-LC (Thermo Scientific, Bremen, GmbH). The peptides were loaded onto a trap column (2 cm × 75 μm) packed in-house using C18 material (Magic C18 AQ, 5 μm, 100 Å) with a flow rate of 3 μl/min using 0.1% formic acid and separated on an analytical column (10 cm × 75 μm, Magic C18 AQ 3 μm, 120 Å) with a flow rate of 350nL/min with a linear gradient of 5% to 60% solvent B (90% acetonitrile, 0.1% formic acid) over a period of 30 minutes. Precursor ions were acquired in m/z range of 350–1800 with a resolution of 60,000 at 400 m/z on an Orbitrap mass analyzer. Ten most abundant ions in every precursor scan were selected and fragmented in HCD mode with normalized collision energy of 39. The dependent scans were performed at a resolution of 15,000 at m/z of 400. Other parameters included capillary voltage of 2.2 kV, capillary temperature was maintained at 250 °C and lock mass option was enabled with polysiloxane ion (m/z, 445.120025) for internal mass calibration.

### Database search

*Mass spectrometry data was* analyzed using SEQUEST search algorithm against *Plasmodium falciparum* 3D7 protein database (version 4.0 containing 5431 protein entries) and frequently observed contaminants through the Proteome Discoverer platform (version 1.3, Thermo Scientific, Bremen, GmbH). The search parameters included a maximum of one missed cleavage; carbamidomethylation at cysteine as a fixed modification; oxidation at methionine, as variable modifications. The MS tolerance was set at 20 ppm and MS/MS tolerance to 0.1 Da. False discovery rate of 1% was applied as cut-off value for reporting identified peptides. False discovery rate was calculated by searching the MS/MS data against decoy database that contained reverse sequences of all proteins in target database. Peptides that passed 1% FDR cut-off were considered as identified. The raw data obtained from these LC-MS/MS runs were submitted to Proteome Commons. Data submitted to PRIDE database refer PRIDE ID 58557.

### Selection of patients

All the patients selected for the study were adults (above 20yr of age). Blood smears, collected from the General Medicine wards of a government tertiary health care facility located in Mangalore, were microscopically examined for malaria parasite from July 2010 to December 2013. The *Plasmodium falciparum* positive malaria cases which met enrollment criteria and provided written informed consent were enrolled under different malaria categories after microscopically confirming *P.falciparum* positive malaria subjects which were classified according to WHO guidelines^26,27,28^. Ethical clearance from Institutional Human Bioethics and Biosafety Review committee was obtained for using patients samples.

### Microarray sample processing

Peripheral blood samples from 11 severe high parasitemia (HP), 4 mild Low parasitemia (LP) patients and 4 *in vitro* grown parasites at ring stage were collected into PAX gene blood RNA tubes and mRNA was isolated using PAXgene blood RNA kit according to manufacturer’s protocol. Samples were prepared for hybridization using Gene chip 3’ IVT express labeling assay kit according to users manual, which included addition of PolyA RNA control followed by first strand and second strand cDNA sythesis. IVT labeling of cRNA was carried out for 4h at 400C and purified using RNA binding beads. Labeled cRNA was fragmented for 35min at 940C according to 49/64 array format and hybridized on Plasmodium/Anopheles Genome Array for 16h at 450C. Staining and washing steps were performed and chip was analysed on Affymetrix platform. The GeneChip® Plasmodium/Anopheles Genome Array which includes probe sets to over 4,300 *Plasmodium falciparum* transcripts and approximately 14,900 *Anopheles gambiae* transcripts were used for studying gene transcription analysis. *P. falciparum* sequence information for this array was collected primarily from PlasmodB and augmented with sequence information from GenBank® and dbEST.

### In vivo Differential Gene Expression Analysis

Our study consisted of two groups of *in vivo* samples severe malaria (HP) and mild malaria (LP) with 4 *in vitro* control samples which were scanned on the Affymetrix GPL1321. Data sets were normalized using RMA software in the Affymertix package of Bioconductor in the R-statistical computing Environment (https://www.r-project.org/). To obtain differentially expressed gene data sets for HP and LP samples, the limma package of R/Bioconductor (Ritchie et al 2015) were used. The FDR-BH (Benjamini–Hochberg) corrected p-value threshold of <= 0.05 and the fold change threshold of the differentially up regulated genes >=1.5 were selected. We obtained 468 HP and 451 LP differentially expressed up-regulated genes from HP and LP analysis respectively. Gini correlation was applied to predict the potential gene interactions. Network was generated using the software Cytoscape (http://www.cytoscape.org). for 468 up-regulated genes of HP with threshold of correlation>= 0.75. For full experimental details and raw data sets are submitted to GEO, refer to the Gene Expression Omnibus website http://www.ncbi.nlm.nih.gov/geo/query/acc.cgi?acc=GSE72448, accession number GSE72448.

### Network construction and Analysis

To understand and elucidate the complex interactions of ETRAMP14.1 with other differentially expressed genes within the 468 up-regulated genes of HP samples, we constructed the gene co-expression network for these genes using their RMA normalized expression values. These expression data sets were further correlated using Gini correlation^29^ with the correlation threshold >=0.75 and found the Gini correlation to be more effective in predicting the potential interactions as compared to Pearson based correlation. The network was constructed using Cytoscape30 which generated 468 nodes (all up regulated HP genes) and 31964 edges (all interactions among these genes with correlation threshold >=0.75. This full network is not shown here). To identify interactions of PfETRAMP14.1 with the proteins in the immediate neighborhood, proteins that interact directly were selected from the whole network which consisted of 468 nodes/31964 edges for all the upregulated genes. From this, we obtained 97 direct (first neighborhood) interacting proteins with 2627 interactions (additional file 1 shows interactions of PfETRMP14.1 rifin, stevor, PfEMP1s, ribosomal proteins and TRX2, PTP6 etc.). Further smaller sub-network was obtained from this by selecting proteins which prominently interact with PfETRAMP14.1 (Figure B) and this network highlights the direct interaction of pfETRAMP14.1 with the proteins in the immediate neighborhood.

### Construction of recombinant ETRAMP14.1 mutants and Immunofluorescence labeling of *E. coli*

Two types of ETRAMP14.1 mutant were generated. First, transmembrane domain mutant (E14.1TM) was constructed using an overlap extension PCR method. Glycine residues in the TM motif (GxGxGxG) were replaced with threonine and alanine (TxAxAxA). The external primers 5’GGGAATTCCATATGAAAGTTTACTAATATTTTATTC-3’; 5’ CCGCTCGAGAGCTGTTGTGGCTGG-3’and internal primers (TTACTTGTTACGACTGCGGTTGCGCTTGCATTCTAC and GTAGAATGCAAGCGCAACCGCAGTCGTAACAAGTAA) were used. Second mutant was C terminus truncated (E14.1CT). DNA sequence of 93bp coding for 31 amino acid residues of C terminus region of ETRAMP14.1 was removed and amplified using 5’GGGAATTCCATATGAAAGTTTACTAATATTTTATTC-3’and 5’CCC<u>CTCGAG</u>GTAGTAGAATCCAAGACCAACACCAGTAC. The PCR products were purified from a 1% agarose gel using the QIAquick gel extraction kit (Qiagen, Inc.) and cloned in NdeI and XhoI sites of pET-22b (+) vector (Novagen) to obatain C-terminus His tagged protein. Cells were fixed in growth medium with 2.4% formaldehyde and 0.04% glutaraldehyde and washed three times with PBS and resuspended in buffer containing 50mM glucose, 10mM EDTA, 20mMTris-Cl pH 7.5. Cells were incubated with lysozyme for 30 min at RT and were washed twice with PBST followed by blocking with 3% BSA for 1 hour. Cells were incubated with α-ETRAMP14.1 antibodies (1:300 dilution) for one and half hour. Alexafluor goat anti-rabbit 488 secondary antibody (Molecular Probes) was added at 1:1000 dilution in 3% BSA/PBS for 1hr.Cells were washed three times in PBS and mounted in 50% glycerol with 0.1mg/ml of DAPI onto the slides which was previously flamed. The coverslips were then inverted and sealed.

## Results

### Detection of ETRAMP14.1 in *P. falciparum* lysate

The full-length ETRAMP14.1 with His-tag at C-terminus was cloned, expressed and purified using Ni-NTA affinity chromatography to generate antibodies in rabbits (Fig S1). The raised antibodies were checked for their specificity by Western blot analysis with the recombinant protein as well as the *P.falciparum* lysate, which showed bands at ∼20kDa and ∼14kDa respectively (Fig 1A). The identity of the protein picked up by the specific antibodies was confirmed by LC-MS/MS as shown in Fig 1B, with a 51.4% coverage.

**Figure 1.**
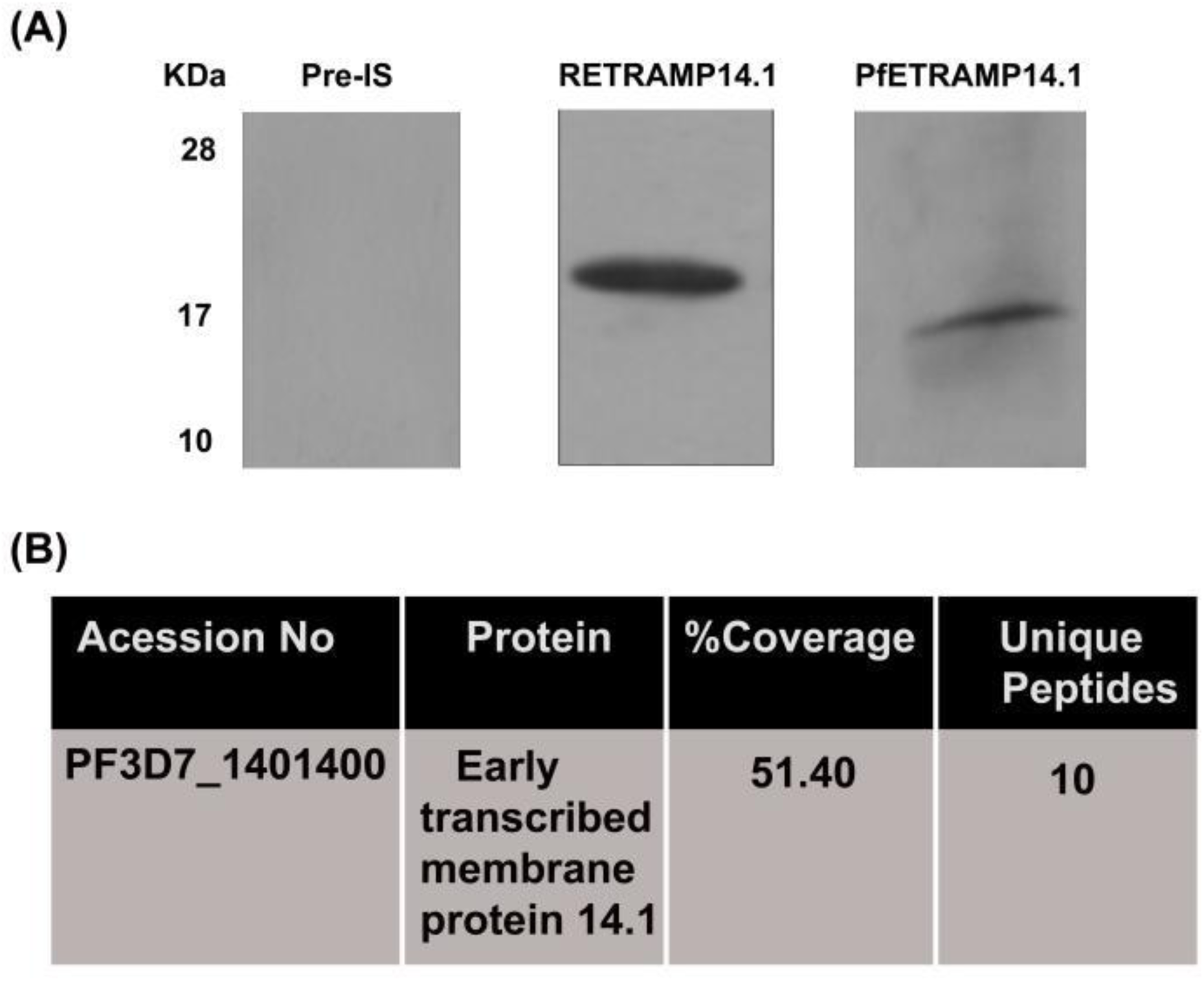
Detection of PfETRAMP14.1 in parasite lysate and confirmation by LC-MS/MS. (A) No signal was detected with pre-immune serum for ETRAMP14.1 (Pre-IS on left), Recombinant protein (REETRAMP14.1-6His) and *P.falciparum* lysate PfETRAMP14.1) probed with anti-ETRAMP14.1 antibodies detected a single specific band at ∼14kDa (right). ETRAMP14.1 shows anomalous mobility as it runs higher than its predicted molecular weight of 11.4kDa. Recombinant protein runs at ∼20kDa due to additional positive charge from His-tag and due to the post translational modifications in *E. coli.* (B) LC-MS/MS identification of PfETRAMP14.1in the *P. falciparum* culture lysate.

### ETRAMP14.1 is expressed in all the intraerythrocytic stages of *P.falciparum*

ETRAMP family of proteins exhibit differential gene expression throughout the life cycle of the parasite. Hence, to understand the transcription and translation pattern of ETRAMP14.1 at different parasite stages, real-time PCR and Western analysis were performed. RNA isolated from rings, early trophozoites, mature trophozoites and schizonts was used for RT-PCR with seryltRNA synthetase as normalization control. The results show that ETRAMP14.1 is maximally expressed in rings (∼30fold increase compared to mature trophozoites, which show lowest expression) whereas the early trophozoites and schizonts show ∼10 fold and ∼5fold increase respectively as compared to mature trophozoites (Fig 2A). We next checked the protein expression levels at different developmental stages while ETRAMP14.1 shows maximum expression in early trophozoites as compared to rings, its expression is lowest in mature trophozoites (Fig 2B). To determine localization of ETRAMP14.1 in different intraerythrocytic stages, synchronized cultures were used for the immunofluorescence assay. ETRAMP14.1 is found to localize to the parasite periphery at PVM/PPM. Strong peripheral signal was seen in rings and early trophozoites as compared to mature trophozoites in which the signal is slightly decreased showing few puncta. Its expression is again increased in the late schizonts showing segmented merozoites. We also observed ETRAMP14.1 expression in mature gametocytes (Fig 2C). Thus, our results indicate that ETRAMP14.1 is expressed during all the intraerythrocytic stages with clear peripheral localization.

**Figure 2.**
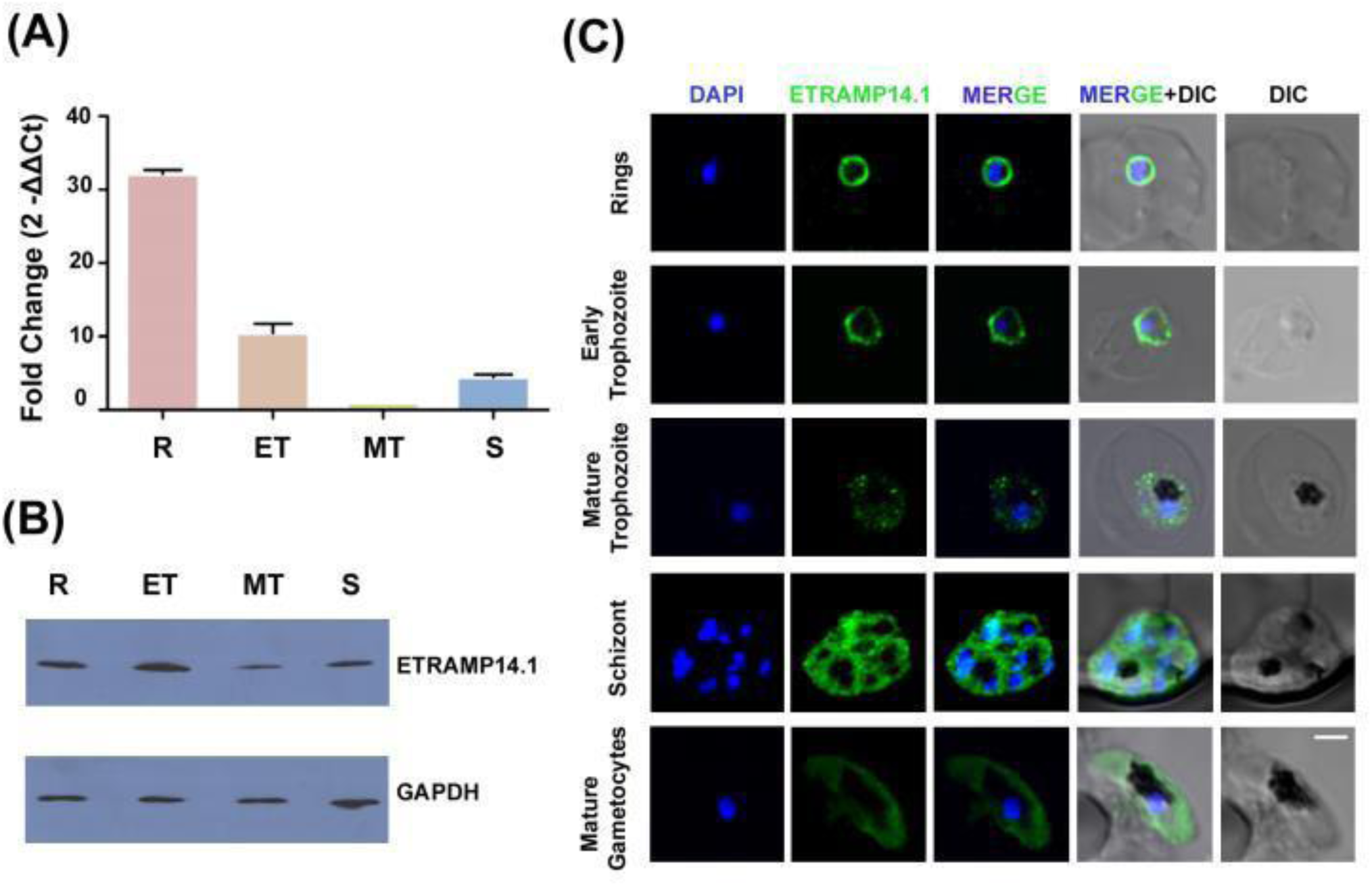
Expression of ETRAMP14.1 at different intraerythrocytic stages (A) Real time PCR analysis of ETRAMP14.1. Transcripts used to amplify ETRAMP14.1were prepared from cDNA obtained at different parasite blood stages; rings (R), early trophozoites (ET), mature trophozoites (MT) and schizonts (S) using Oligo-dT primers. (B)Western blot analysis of ETRAMP14.1 at different intraerythrocytic stages; ring (R), early trophozoites (ET), mature trophozoites (MT) and shizonts (S). GAPDH is used as loading control (C) IFA of ETRAMP14.1 in different intraerythrocytic stages. Scale bar: 2μm.

### ETRAMP14.1 is a PVM resident protein

Western analysis using soluble and insoluble fractions of parasite lysate show association of ETRAMP14.1 with the membrane fraction (Fig 3A). In order to determine the exact localization of ETRAMP14.1, colocalization studies were performed with PVM markers, PTEX150 and HSP101, the components of PTEX. Complete overlap between the ETRAMP14.1 with PTEX150 (Fig 3B) and HSP101 (Fig 3C) at different intraerythrocytic stages of the parasite was found confirming the localization of ETRAMP14.1 at PVM. Furthermore, its PVM localization was confirmed by selective permeabilization. IFA was performed after treatment with Streptolysin O (SLO) which permeabilizes only RBC membrane but not PVM and PPM (antibodies have access to only PVM but not to PPM). Only SLO treated parasites showed ETRAMP14.1 signal which was concentrated at specific sites suggesting ETRAMP14.1 forms discrete domains at PVM (Fig 3D), NUP100 did not show any signal in SLO treated parasites which proves SLO permeabilizes only RBC membrane with no access to nuclear periphery. In positive control, NUP100 showed signal at the nuclear periphery when iRBCs were treated with TritonX-100 which permeabilizes all the membranes (Fig S2). ETRAMPs have unique C-terminus region and is speculated to be required for PVM localization^8^. C-terminus deletion mutant of ETRAMP14.1 generated in our transfection studies suggests that C-terminus is not required for PVM localization (data not shown). IFA was also performed with *E. coli* cultures expressing all the three recombinant forms i.e.,; full length (E14.1FT), transmembrane domain (E14.1TM) mutant and C-terminus (E14.1CT) deletion mutant as shown in Fig S3. All of them showed membrane localization suggesting that ETRAMP14.1 can self-associate in membrane without the aid of other *Plasmodium* proteins.

**Figure 3.**
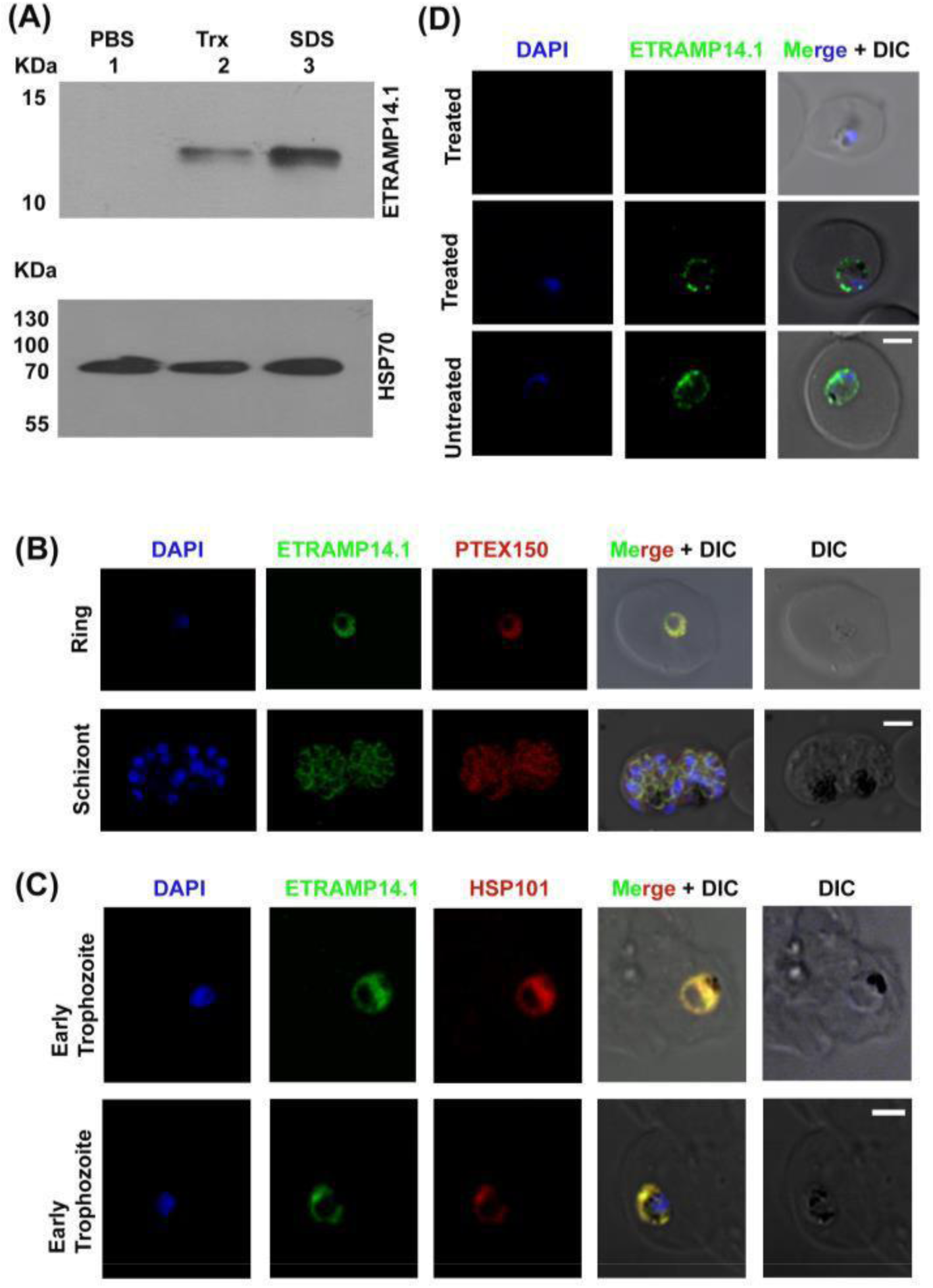
ETRAMP14.1 localizes to parasitophorous vacuolar membrane (PVM). (A) Western analysis of soluble (lane1-parasite protein extracted with PBS) and insoluble fractions (lane 2 and lane 3-parasite proteins extracted with TritonX-100 and SDS respectively) of parasite proteins probed with anti-ETRAMP14.1. Hsp70-1 is used as loading control (B) & (C) Colocalization of ETRAMP14.1 with PTEX150 and HSP101 respectively. (D) IFA of selectively permeabilized iRBC. Upper two panels show IRBCs permeabilized with SLO, enabling access of antibodies through the permeabilized RBCs resulting in fluorescence signal if the corresponding antigen is located in the PVM, while the bottom panel is control IRBCs without SLO treatment showing no signal. Scale bar: 2μm.

### IP-LC-MS/MS analysis reveals interaction of ETRAMP14.1 with PfEMP1 andits other interacting partners

In an attempt to identify the interacting partners of ETRAMP14.1, which might help in understanding its functions, immunoprecipitation of the total parasite lysate with ETRAMP14.1 antibodie was carried out (Figure 4A). IP-LC-MS/MS analysis identified a total of 12 proteins (Figure 4B) which includes Hsp70-1, EXP-2, PHISTc, 14-3-3 and PfEMP1. PfHsp70-1 was detected in IP at 1% FDR with one unique peptide where as 14 of its peptides were identified at 5% FDR. EXP2 was identified with one unique peptide SHTLYTHITPDAVPQLPK at 1% FDR whereas three other unique peptides were detected at 5% FDR. Similarly, PfEMP1 was detected with one unique peptide (AITCHVVSGNNYFR) at 1% FDR and 3 other peptides (AITCHVVSGNNYFR, IRTIDDDLIK, EYWWNANR) at 5%FDR. Other proteins found in LC-MS/MS analysis are listed in Figure 5C. Refer PRIDE ID 58557 for full data. The results obtained in LC-MS/MS analyses were validated by probing Western blots of immunoprecipitated complex (pulled down with anti-ETRAMP14.1 antibodies) with anti-Hsp70-1 and anti-PfEMP1 antibodies (Figure 5A & 5B). Both the proteins gave signals at the expected molecular weight of 73kDa and 250kDa respectively. These results further confirm the association of ETRAMP14.1 with the chaperone Hsp70-1 and PfEMP1.

**Figure 4.**
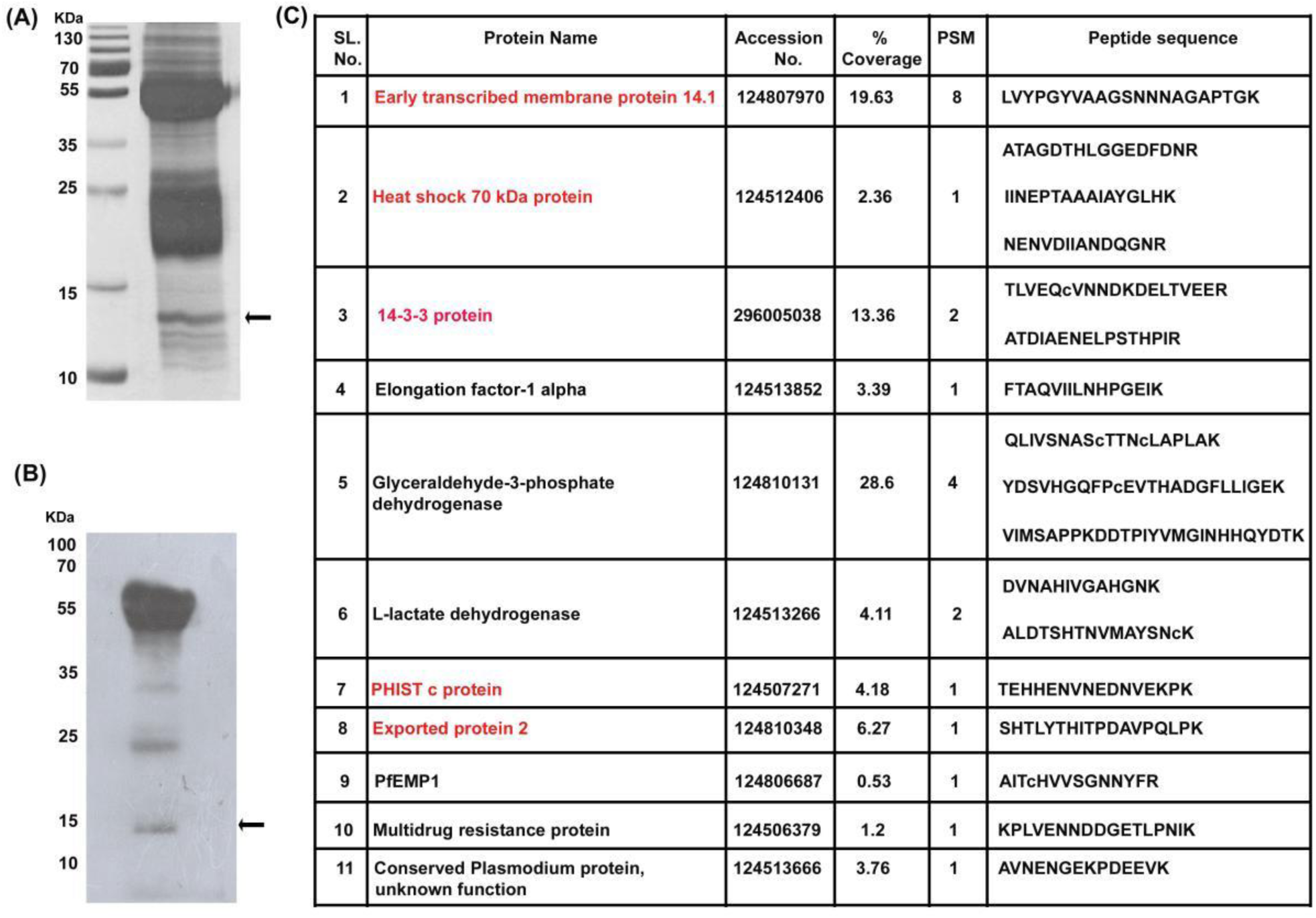
Identification of ETRAMP14.1 interacting partners by LC-MS/MS on immunoprecipitated complex. (A) Silver stained gel of immunoprecipitated complex. (B) Western blot probed with anti-ETRAMP14.1 (C) List of interacting partners of ETRAMP14.1 identified by LTQ XL mass spectrometer. Raw data files were searched using Proteome Discoverer 1.3 and the SEQUEST algorithm against the database. Proteins highlighted in red are found in parasitphorous vacuole and PVM, among which some are present even in parasite cyotosol. Arrows indicate ETRAMP14.1 in the IP complex.

**Figure 5.**
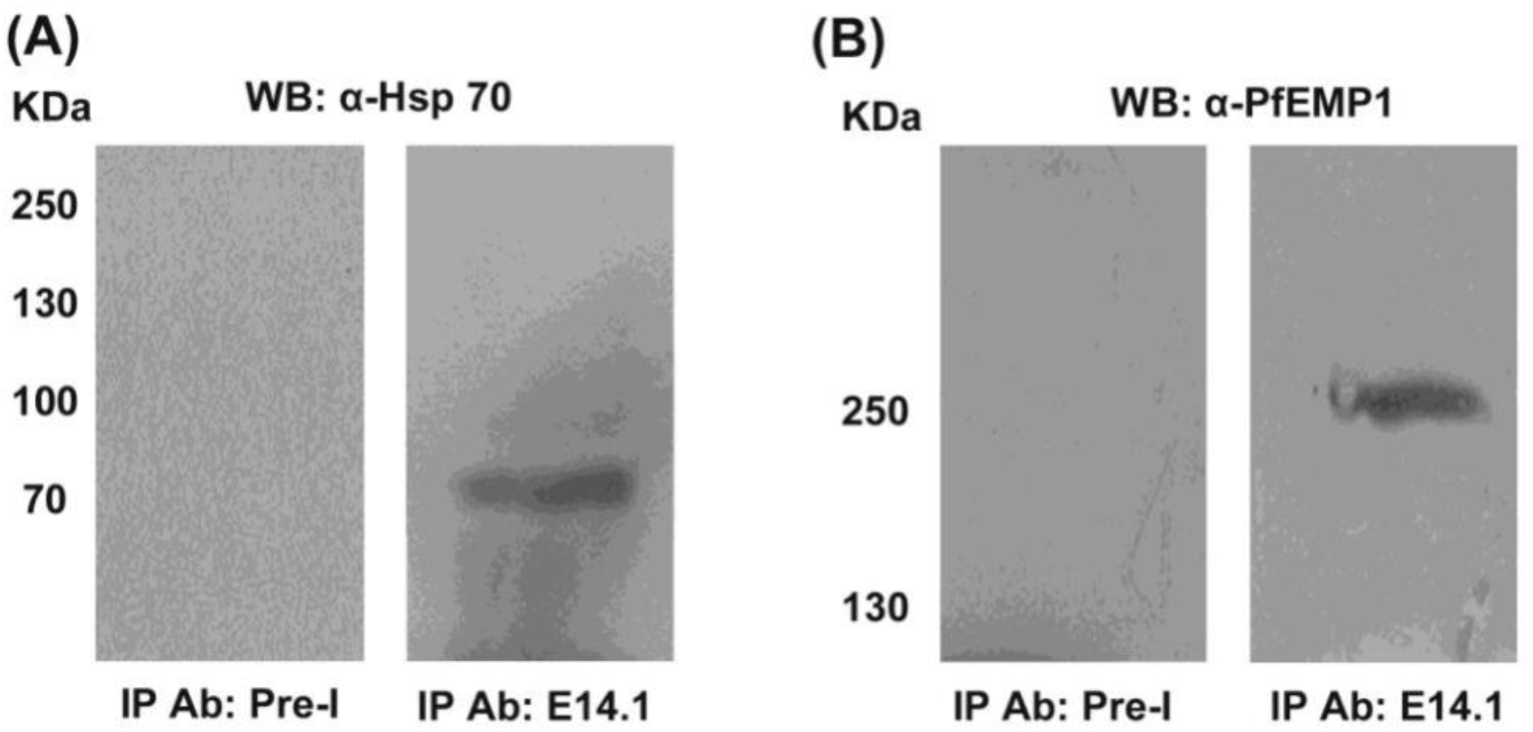
ETRAMP14.1 interacts with PfHsp70-1 and PfEMP1 (A) Western blot of immuno-precipitated complex pulled down with pre-immune serum (left) and anti-ETRAMP14.1 (right) antibodies, with anti-Hsp70-1. (B) Western blot of immuno-precipitated complex pulled down with pre-immune serum (left) and anti-ETRAMP14.1 (right) probed with anti-PfEMP1.

### Reconstructed protein-interaction network shows PfEMP1, a PTEX protein and few other ETRAMP members as its interacting partners corroborating the *in-vitro* results

As, the *in vitro* proteomic analyses show PfEMP1 and EXP2 (a PTEX component) as interacting partners of PfETRAMP14.1, we were fascinated to know whether these interactions are a true reflection of the *in vivo* scenario. For this, array analyses were carried out on parasite RNA isolated from peripheral blood samples of 11 severe malaria (HP, high parasitemia) and 4 mild malaria (LP, low parasitemia) patients. We found 468 and 491 genes to be up-regulated in HP and LP samples respectively with upregulation of 241 genes in common. The 25 most commonly upregulated genes in severe malaria are represented in the bar graph (Figure 6A). Genes such as PfEMP1, PHISTc, ETRAMP14.1, ETRAMP5, ETRAMP11.2, ETRAMP10.1, PTP6, TRX2, and 14-3-3 are upregulated in high parasitemia samples as shown in Figure 6A (subpanel). Among these genes, interestingly, ETRAMP14.1 is maximally expressed (∼40fold upregulation) *in vivo* in severe malaria patients. To understand the interactions of ETRAMP14.1 with other differentially expressed genes within the upregulated genes of severe malaria samples, a protein interaction network was reconstructed using RMA normalized expression data of the 468 up regulated genes of HP samples. Based on this network analyses, we identified a number of interacting partners of ETRAMP14.1, such as surface adhesins (PfEMP1s, RIFINS and Stevors), PVM proteins, ribosomal proteins etc. (Figure S4). A subnetwork showed direct interactions of ETRAMP14.1 with selected genes such as PfEMP1, ETRAMP11.2, ETRAMP10.1, TRX2, PTP6, Plasmepsin VII and 14-3-3 (Figure 6B). Thus, it is clear that PfETRAMP14.1 does interact with PfEMP1 and translocon machinery. Large network clusters around ETRAMP14.1 are shown in Figure S4.

**Figure 6.**
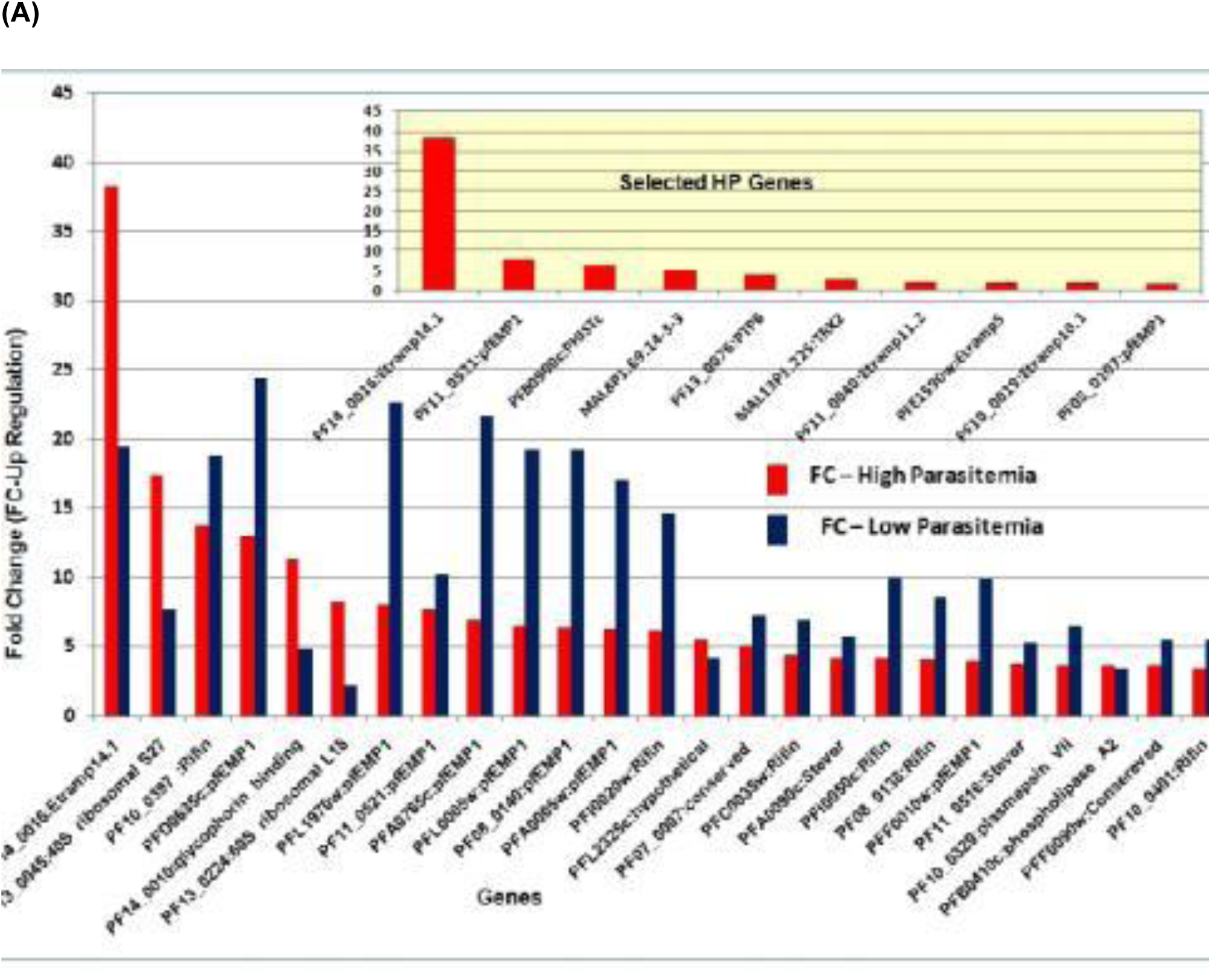
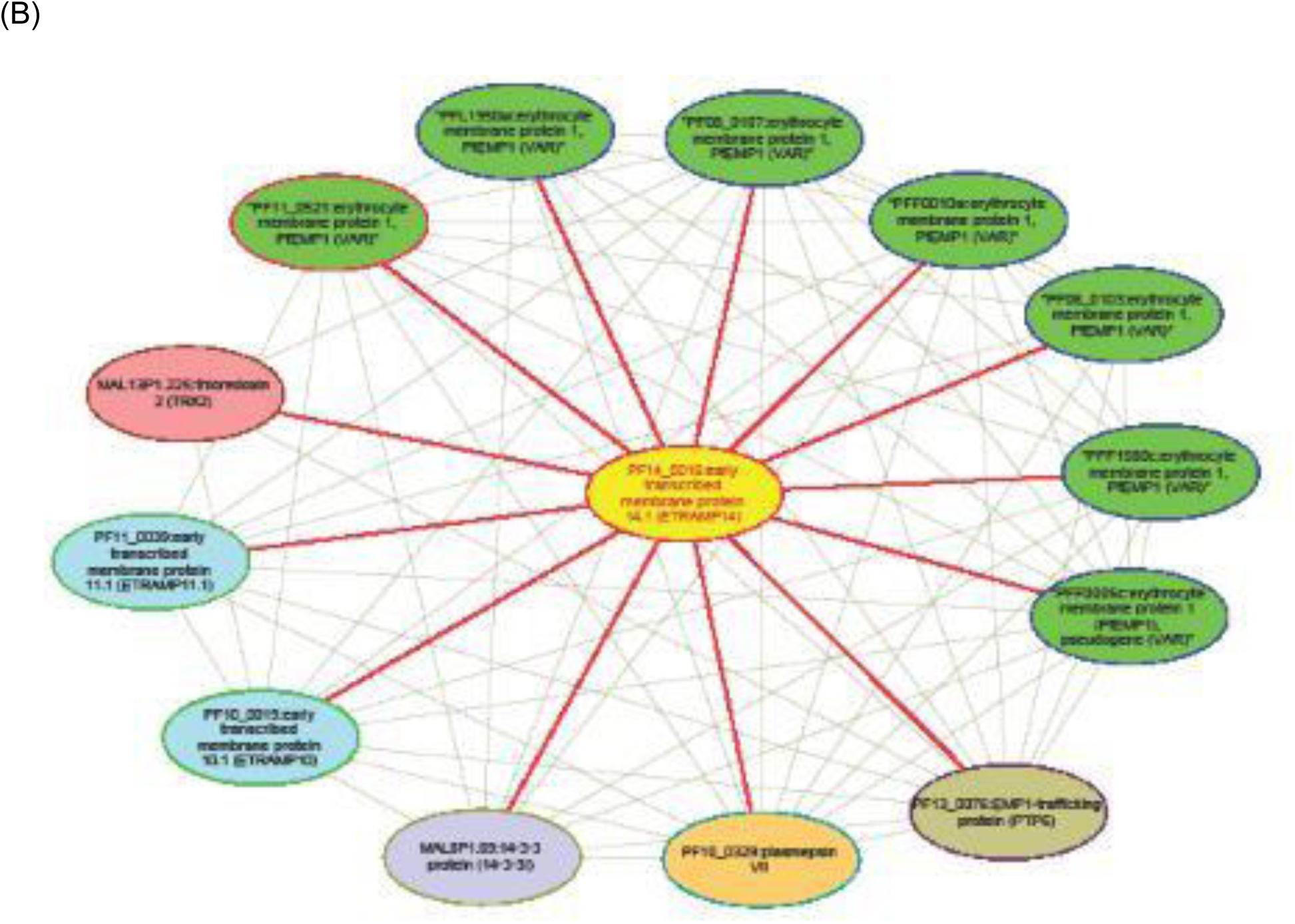
*In vivo* expression of ETRAMP14.1 and reconstructed ETRAMP14.1 protein-interaction sub network (A)Comparison of *in vivo* fold change (FC) for top 25 genes showing highest expression in *Plasmodium falciparum* samples of severe (HP) and mild (LP) malaria samples. The sub panel highlights the upregulation of some of the selected genes from the severe samples such as ETRAMP14.1, PfEMP1, PHISTc, ETRAMP5, ETRAMP11.2, ETRAMP10.1, PTP6, TRX2, and 14-3-3. Fold change threshold: up =(+1.5). (B) Subnetwork of *in vivo* ETRAMP14.1-interacting proteins. The different color nodes represent genes while solid lines represent interactions. Gene names are as given in PlasmoDB. ETRAMP14.1 (yellow), PfEMP1 (green) Thioredoxin 2 (red), two ETRAMP members ETRAMP11.1 and ETRAMP10.1 (blue), 14-3-3 (purple), Plasmepsin VII (orange) and PTP6 (brown). Solid red lines show direct protein-protein interactions of ETRAMP14.1 with other genes while the grey lines show indirect protein–protein interactions. This protein-protein interaction network of ETRAMP14.1 predicts PfEMP1s, PVM proteins and other ETRAMP members as interacting partners of ETRAMP14.1. Additional partners are shown in a larger network in Figure S4.

### A proposed model for ETRAMP14.1 function

Based on our proteomic studies and reconstructed protein interaction network, we propose a model for the interactions of ETRAMP14.1 with its partners. Discrete domains of ETRAMP14.1 as demonstrated by selective permeabilization studies and its potential interaction with parasite proteins such as EXP2, PfHsp70-1, TRX2, PfEMP1 and other ETRAMP members are represented in Figure 7. We speculate that PfHsp70-1 being a chaperone, helps in proper folding of ETRAMP14.1 and might stabilize its localization at PVM or it may even help in exporting ETRAMP14.1 to PVM. PTEX is a macromolecular complex with EXP2, forming the core channel for proteins to be exported into host cytosol to host RBC surface, a crucial process essential for virulence and viability of parasite. PfEMP1 or any of the exported protein needs to be unfolded and delivered through the translocon machinery. We propose that PfHsp70-1 might aid in export of PfEMP1 or regulate EXP2 of the translocon to facilitate trafficking of PfEMP1 and other proteins to host iRBC cytosol and surface.

**Figure 7.**
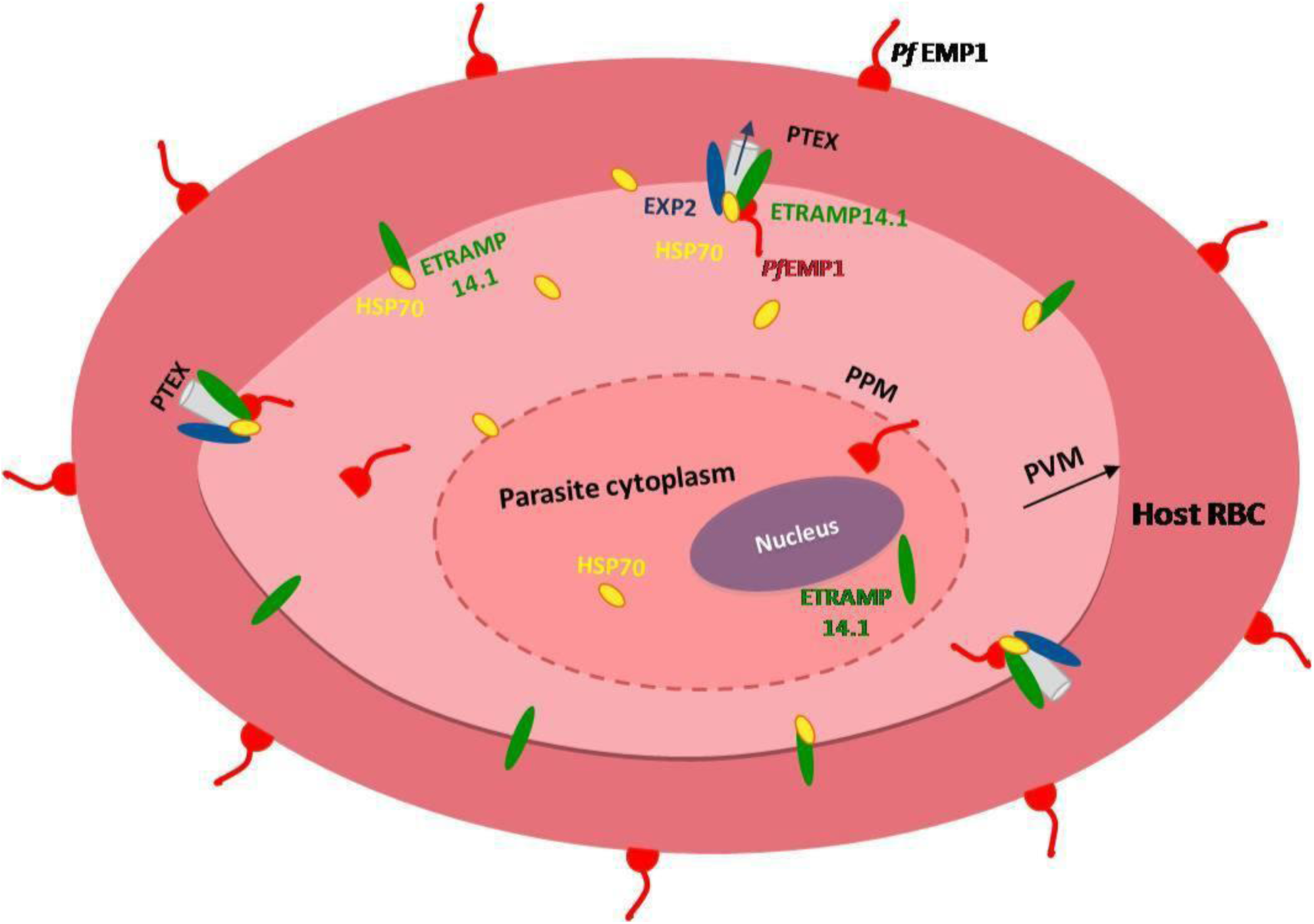
A model for ETRAMP14.1 interaction at PVM. ETRAMP14.1 localizes to PVM forming subdomains and interacts with parasite proteins such as PfEMP1, Hsp70-1, TRX2, EXP2 (a core component of PTEX export machinery) and other ETRAMPs, ETRAMP10.1 and ETRAMP11.1.

## Discussion

Our study provides experimental evidences for the identification of *in vivo* as well as *in vitro* interacting partners of ETRAMP14.1, thus delving for the first time insights into the interactome of ETRAMP14.1. The interactions in the reconstructed in vivo protein interaction network on the array data obtained from severe malaria inflicted patients were validated by in vitro IP-LC-MS/MS analyses identifies PfAMP1, PTEX components, Hsp70-1 and other ETRAMP members as interacting partners of ETRAMP14.1. In addition, PTP6 and PHISTc, the known PfEMP-1 interacting partners too were identified.

Importantly ETRAMPs including ETRAMP14.1 are reported to be recognized by sera of naturally exposed individuals^10^. The fact that ETRAMP14.1 is found to be the highest transcribed gene in severe malaria patients from a malaria endemic region of India, its potential as a biomarker for the specific diagnosis of *P. falciparum* infection needs to be explored. Earlier studies have reported ETRAMP14.1 to bearing stage specific^10^, but in our studies we observed its expression throughout all the intraerythrocytic stages. However, it is maximally expressed in rings through early trophozoites, indicating that it might be involved in the host RBC modifications which takes place during early parasite development, essential for parasite survival. Further, ETRAMP14.1 is minimally expressed in mature trophozoites indicating that it might be replaced by other ETRAMPs to cater stage specific developmental needs of the parasite. ETRAMP14.1 is also expressed in mature gametocytes suggesting that it might have a role in development of sexual stages too. Some ETRAMP memberssuch as ETRAMP2, ETRAMP10.1, and ETRAMP4 form microdomains in PVM as demonstrated earlier by *in vivo* cross-linking studies^11^. In our selective permeabilization studies, we also observed bead like fluorescence for ETRAMP14.1 distributed unevenly throughout the PVM indicating that it is forming microdomains.

Our *in vivo* and *in vitro*protein interaction data also emphasize the interaction of ETRAMP14.1 with other ETRAMP members ETRAMP10.1, ETRAMP11.2 and ETRAMP5 which might be required to perform its functions. Also this may be essential for ETRAMP family proteins to stably assemble in PVM which corroborates with *in vivo* cross-linking studies reported earlier^11^.

PfHsp70-1 has been reported to be present on parasitophorous vacuole^31^ to some extent apart from its cytosolic location. We speculate that it might interact with ETRAMP14.1 to aid proper folding or loading of ETRAMP14.1 onto PVM. The major virulent protein, PfEMP1is exported to host RBC surface within 20 hours post invasion^32,33,20^ and thereby all the translocon components and necessary proteins are required to be expressed much earlier to facilitate the export. ETRAMP14.1 protein expression *in vitro* is found to be maximum at 0-30h post invasion corresponding to the timing of PfEMP1 export. Altogether, these observations indicate that ETRAMP14.1 might be involved in trafficking of PfEMP1 through PVM with the help of EXP2. We also speculate that it might be a component of the translocon, which is a macromolecular complex, as ETRAMP14.1 is present at PVM and shows interaction with EXP2 in our studies.

It is worth mentioning that in our proteomic analyses, PHISTc was also detected as one of the interacting partners of ETRAMP14.1. The central flexible segment of ATS of PfEMP1 is known to associate with PHISTc (PFI1780w) required for cytoadherence of PfEMP1^29^ and this could be the reason for co-immunoprecipitation of PHISTc with PfEMP1. Altogether, this study reveals the participation of an ETRAMP14.1 in trafficking the major virulent protein, PfEMP1 to the surface of host RBC in association with translocon components EXP2 and TRX2; the chaperone HSP70-1 and other ETRAMP members to perform its functions. Hence we speculate that ETRAMP14.1 possibly requires the chaperone Hsp70-1 for proper folding during its export to PVM and facilitates the export of PfEMP1 /other parasite proteins from PVM via the translocon machinery to the host RBC surface. Understanding the mechanism of assembly and various functions of this fascinating family of ETRAMPs will pave way for dissecting trafficking of ‘Var’ antigens to host cell surface and also in developing novel antimalarials.

## Acknowledgements

The authors thank late Dr. Neeru Singh, RMRCT-Jabalpur, Dr. S. K Ghosh and Dr. Pradeep, NIMR, Bangalore for providing clinical isolates of malaria. We thank Bindu M. and Lavanya T. for RNA isolation and array hybridization of patient samples, we thank Dr. Shipra from BiocosLife Sciences for microarray data and protein network analysis, Prof. Utpal Tatu to provide Hsp70-1 and PfEMP1-ATS antibodies as kind gift. We also thank Anitha G. for scanning the gene chip and Suma B.S. for acquiring confocal images. We gratefully acknowledge Keshava Datta, Keshava Prasad and Akhilesh Pandey at The Instituite of Bioinformatics, ITPB, Bangalore for the proteomics analyses. The funding for the project Indian Council of Medical Resarch (ICMR). NS also acknowledges partial support from the JNC-DBT partnership programme (BT/INF/22/SP27679/2018).

## Authors’ contributions

N.S and K.M.V conceptualized the study. K.M.V conducted the experiments and wrote the manuscript. N.S and A.R corrected the manuscript and compiled the data. S.K performed micro array data analysis and generated Protein-Protein interaction network. All authors read and approved the final manuscript.

