## Supplementary Figures for "Interactions of *Plasmodium falciparum* ETRAMP14.1 with PfEMP1, translocon components and other ETRAMP members at the interface of host-parasite"

**Figure S1**

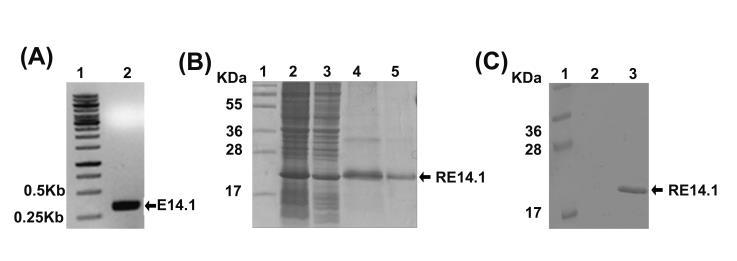

Cloning, expression and purification of recombinant ETRAMP14.1 (A) PCR amplification of ETRAMP14.1

(B) Overexpressed and purified recombinant PfETRAMP14.1-His tagged protein visualized in a coomassie-

stained SDS-PAGE gel. (C) Western blotting of recombinant ETRAMP14.1 with anti-His antibodies.

**Figure S2**

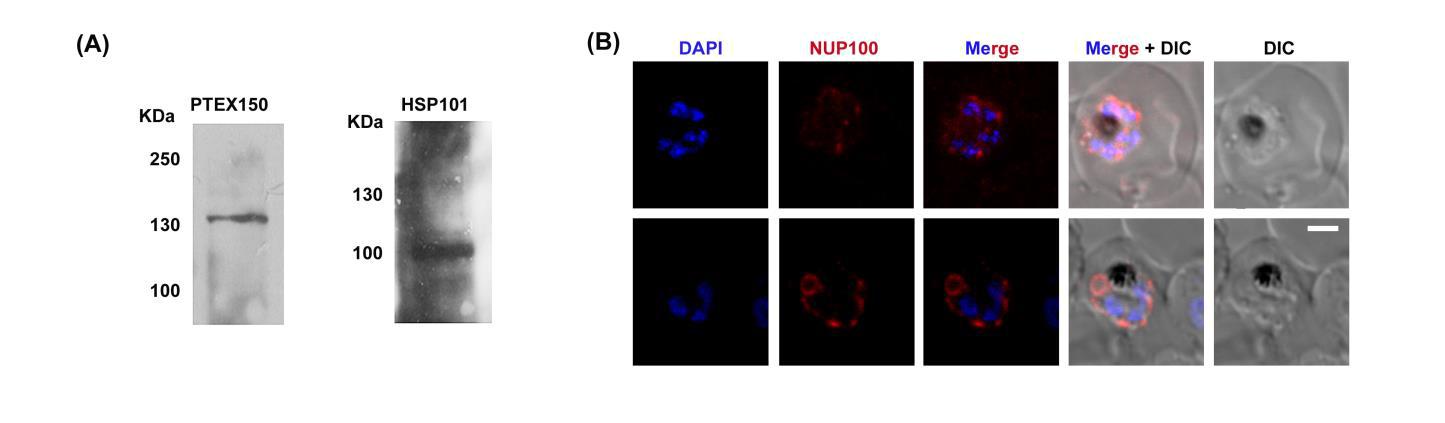

Specificity of α-PTEX150 and α-Hsp101 antibodies and IFA of Nup100. (A) Western blot analysis of

parasite lysate probed with α-PTEX150 (left) and α-Hsp101 (right) antibodies. (B) Immunofluorescence

analysis of infected eythrocytes using Nup100 in TritonX-100 permeabilized cells (used as control showing

TrionX-100 permeabilizes all the membranes; RBC membrane, PVM and PPM).

**Figure S3**

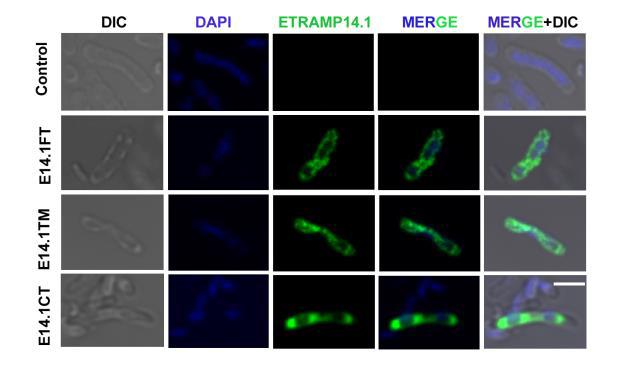

IFA of recombinant ETRAMP14.1 in *E. coli* expressing wild type (E14.1FT) and mutant (E14.1TM,

transmembrane domain mutant and E14.1CT, C-terminus truncated) proteins.

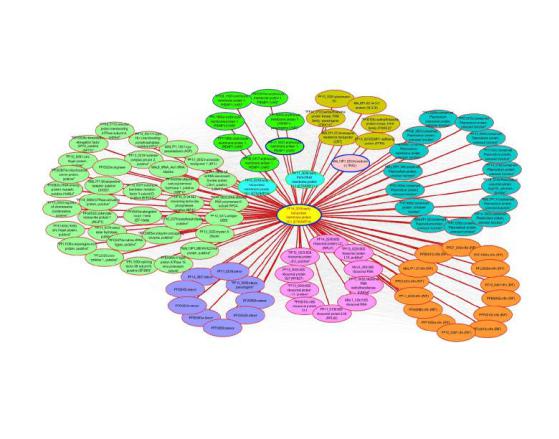
**Figure S4**

Network clusters around PfETRAMP14.1.

26
